# Exploring Neural Signatures of Self-Initiated Saccadic and Blink Events in the Real World

**DOI:** 10.64898/2026.07.13.737665

**Authors:** Aitana Grasso-Cladera, Debora Nolte, Alina Zaidan, Aziz Muhammed Akkaya, Tim C. Kietzmann, Peter König

## Abstract

During natural vision, the human brain constantly has to account for self-produced interruptions of visual input, through blinks and saccades, to maintain a stable perception of the world. It remains unclear whether these event-related neural responses are treated similarly due to a common underlying mechanism. To systematically investigate this functional connection, we recorded synchronized mobile EEG and eye-tracking data from freely moving participants as they (visually) explored a city center. Our results replicate prior findings showing that a substantial portion of the neural response is aligned to event onsets in comparison to offset, underscoring the relevance of investigating the beginning and end of these events. Using advanced deconvolution techniques, we disentangled overlapping, time-locked neural contributions associated with the onsets and offsets of saccades and blinks. The results demonstrate that the deconvoluted neural kernels are highly correlated when comparing saccade onset to blink onset, as well as saccade offset to blink offset, for both evoked and induced activity across electrodes. Together, our findings demonstrate that despite their distinct physical profiles, saccades and blinks share a common neural signature that governs both their onset and offset dynamics during unconstrained real-world behavior.

**Highlights:**

- Co-registration of portable EEG and Eye-Tracking yielded meaningful real world data.
- We studied blinks and saccades as natural interruptions of visual processes.
- Aligning EEG data to onset and offset underscored the relevance of both timepoints.
- We deconvoluted overlapping neural activity for onsets/offsets of saccades/blinks.
- ERPs and TFRs are highly correlated for onset/offset of blinks and saccades.

## Introduction

Natural vision arises from the continuous interaction between perception and action, with sensory input being actively shaped by self-generated movements (O’Regan & Noë, 2001), including naturally occurring eye events that alter visual input, such as saccades and blinks. Saccades and blinks are brief interruptions of the visual stream, with one shifting the gaze while the other occludes the visual field through the closing of the upper eyelid. Despite their difference in the disruption and subsequent continuation of the visual input, these events are both generally unperceivable, revealing consequences indicative of a mechanism that maintains perceptual continuity. Indeed, saccades and blinks are accompanied by visual suppression, commonly interpreted as suppression of blurred visual input during fast eye movements or eyelid closure (Burr, 2005; Johns et al., 2009; Ridder & Tomlinson, 1997; Willett et al., 2023), though it is unclear whether the underlying mechanisms of these suppressions are shared (Willett et al., 2023). Despite their distinct signatures, perception during saccades and blinks is stabilized by similar mechanisms that preserve perceptual continuity and prevent disruptions to the visual experience.

Previous studies have investigated neuronal processes associated with saccades and blinks. In humans and non-human primates, blink- and saccade-related activity involves overlapping brain areas, suggesting shared involvement of oculomotor and visual processing networks (Bodis-Wollner et al., 1999; Shibutani et al., 1984). Event-related studies in humans have further shown that both saccades and blinks elicit reliable, structured neural responses, indicating cortical processing around these self-generated interruptions to visual input (Jagla et al., 2007; Wascher et al., 2014; Wunderlich & Gramann, 2021; Yang et al., 2024). ECoG recordings in humans additionally revealed comparable gamma-band modulation patterns in the early visual cortex for both blinks and saccades, with decreases in gamma power at event onset and increases at event offset (Kern et al., 2021). Extending this temporal perspective by using free viewing to investigate event-related potentials, recent work has similarly demonstrated that both saccade onsets and offsets play different but important roles in the segmentation and processing of visual input (Amme et al., 2024; Nolte et al., 2025). Together, these findings suggest a close functional relationship between blink-and saccade-related processes and highlight the contribution of onset- and offset-related dynamics to the processing of self-generated visual interruptions.

The role of event onsets and offsets in structuring visual processing raises the question of whether these dynamics generalize to natural vision during real-world exploration, where visual input is continuously shaped by interactions between perception and action. Additionally, it remains unclear whether blink-related processing is similarly shaped by contributions from onset and offset. Investigating blink- and saccade-related neural signatures during unconstrained real-world behavior may provide insight into how these dynamics compare across events.

The current study systematically investigated this question by recording synchronized mobile EEG and eye-tracking data from freely moving participants as they explored the city center of Osnabrück, Germany. Using the eye-tracking data to identify relevant time points, we compared neural signatures across different eye events. We explored the relevance of saccade and blink onsets and offsets, aiming to replicate previous work (Amme et al., 2024; Nolte et al., 2025), and hypothesized that saccade onset would better explain early ERP components, such as the P100 component, than saccade offset (i.e., fixation onset), and extended these results to blink events. By performing a within-subject comparison using deconvolution techniques for the onsets and offsets of saccade and blink events, we aimed to disentangle the distinct neural contributions of these event onsets and offsets. Overall, this study seeks to contribute to understanding how the human brain sequentially orchestrates visual suppression and perceptual updating during unconstrained real-world exploration.

## Methods

### Participants

A total of 28 subjects participated in the experiment, resulting in 56 sessions (2 per subject). To prioritize high-quality data, 28 of these sessions were excluded from the analysis: both sessions from 6 subjects and single sessions from 2 additional participants were discarded due to technical issues during recording. Further, both sessions from 5 other subjects and single sessions from 4 other subjects were discarded due to high noise level in the EEG recordings (N = 10) or absence of one of the eye-tracking markers for data alignment (N = 2). This resulted in a total of 28 sessions (16 subjects with both sessions and 4 with a single session) included in the final dataset (9 female, zero diverse; mean age 27 ± 4.63). Fifteen of the subjects reported being right-handed, and the majority of them (N = 12) were bachelor’s students or had completed the bachelor’s level. All included subjects had normal or corrected-to-normal vision, did not report any neurological diagnoses, provided written informed consent before participation, and were compensated with participation hours or a monetary reward, as preferred by the participant. The present study was conducted in accordance with the Declaration of Helsinki and was approved by the University of Osnabrück’s ethics committee.

### Apparatus

We recorded eye tracking data from both eyes and the world camera using the Pupil Lab Neon glasses (Pupil Labs, Berlin, Germany; first panel, Figure 1A). The eye-tracker recorded gaze positions at 200 Hz, a resolution of 1600 x 1200 pixels, and an uncalibrated accuracy of 1.8° (Baumann & Dierkes, 2023). Moreover, we recorded and analyzed the video from the world camera (RGB) with a sampling rate of 30 Hz, a 132° x 81° field of view, and a resolution of 1600 x 1200 pixels. Data was recorded locally through the Companion App (Pupil Labs, Berlin, Germany) on a Samsung S23+ smartphone and uploaded to PupilCloud (Pupil Labs, Berlin, Germany). We computed the effective sampling rate for both the world and eye cameras to validate their consistency over time (Supplementary Material 1; see the *Validation of the Eye-Tracking Data* section).

**Figure 1.**
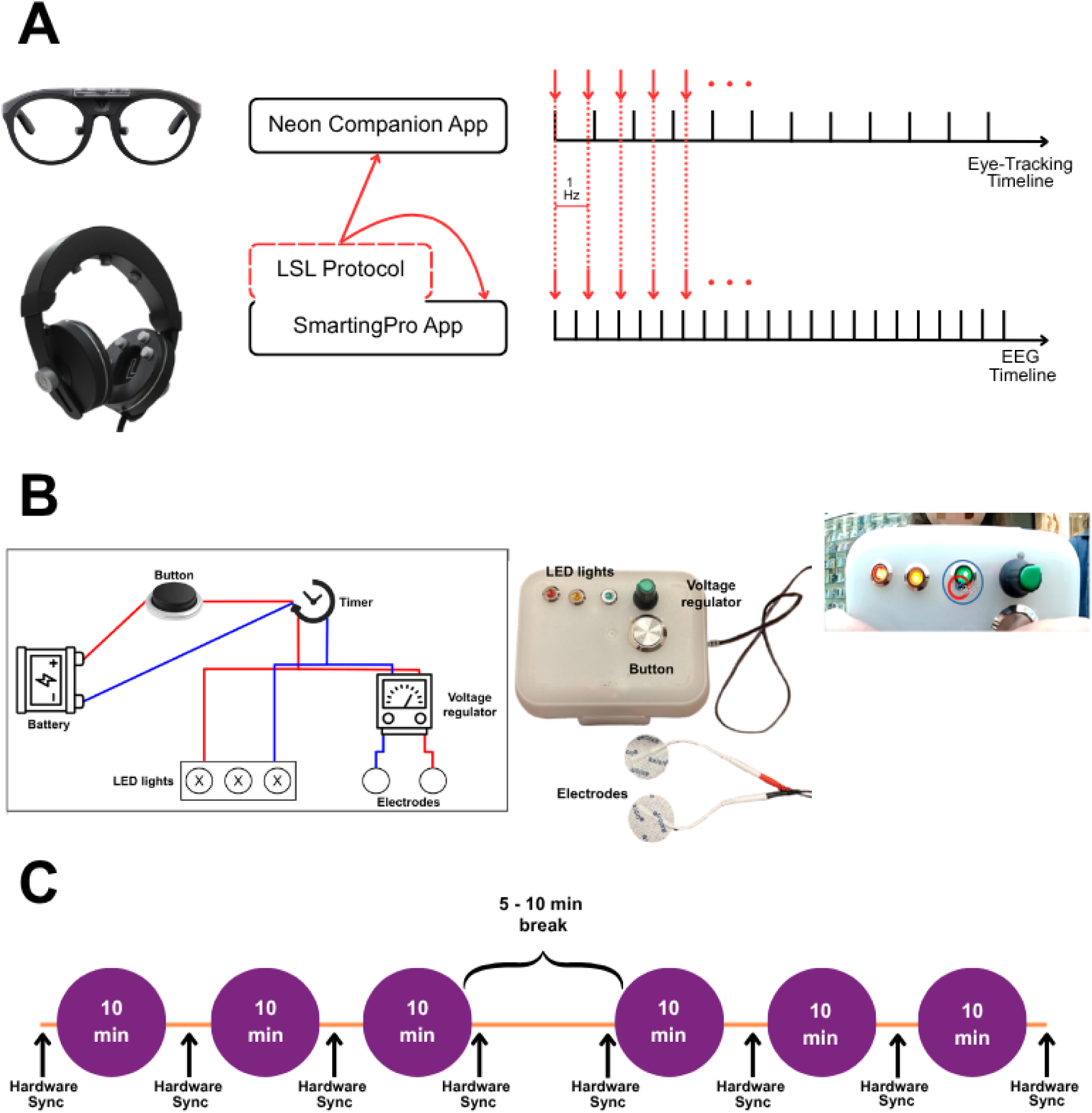
Synchronization protocol at the software and hardware levels. (A) illustrates software-level synchronization, in which the EEG APP (SmartingPro) sends triggers at 1-second intervals to both the EEG and eye-tracking data streams. (B) depicts the hardware-level setup, which involves a circuit designed to generate simultaneous artifacts in both streams via visual signals on the eye tracker’s world camera and electrical signals on the EEG by placing two electrodes on the participant’s face near the EEG headset. (C) Illustrates the general structure of the experiment, which consists of two 30-minute sessions of data recording during free exploration, followed by a 5- to 10-minute break, and the generation of 8 rounds of hardware-level artifacts. (D) Displays the location of the experiment. The pictures are property of the authors and the map source is Google.

For recording EEG data, we used the Smarfones system (mBrainTrain; Belgrade, Serbia; first panel, Figure 1A) equipped with 11 sponge-based data electrodes and reference and ground electrodes incorporated in the headset. We prepared the headset according to the manufacturer’s instructions by soaking the sponges in a saline solution for at least 12 hours before the recording. The data electrodes were referenced to the built-in reference electrode, positioned at the bottom of the left earcup at the left mastoid. The signal was amplified using the Smarting Pro amplifier (mBrainTrain; Belgrade, Serbia), and data were recorded at a sampling rate of 500 Hz. The amplifier was connected via Bluetooth to the SmartingPro App (mBrainTrain; Belgrade, Serbia) on the same Samsung S23+ smartphone, where data was collected and saved locally in an XDF-file.

As precise time information is essential for the current study, we employed a two-step approach to synchronize the eye-tracking and EEG datasets. First, we implemented a software synchronization procedure using an HTTP protocol developed by mBrainTrain, through which the SmartingPro App sent triggers to the Companion App (Jovanovic, 2024). We used the *RESTAPI* LabStreamLayer (LSL; (Kothe, 2014; Kothe et al., 2024) stream to send triggers every second while recording, ensuring precise synchronization (middle and right panels; Figure 1A). Secondly, we used a hardware synchronization validation routine (Figure 1B). For this, we introduced simultaneous physical artifacts in the eye-tracking and EEG signals by presenting LED lights in front of the eye-tracker’s world camera and by sending a simultaneous ∼2.5 V impulse to the EEG headset through two electrodes positioned on the participant’s face (Figure 1B). As a sanity check, we calculated the difference between the light and electrical impulse generation to control for any potential temporal shift introduced by the electrode voltage regulator. As expected, we identified a small constant offset, with a mean of 1.122 ms (median = 2 ms; standard deviation = 1.004 ms), indicating that the light tends to appear faster than the electrical impulse. We accounted for this drift in the data alignment procedure (see the *Data Alignment* subsection in *Preprocessing*). During each session, the hardware synchronization procedure was performed every 10 minutes of recording, and for each of these, we sent 10 consecutive artifacts, unevenly spaced, to ensure device synchronization in case of linear or nonlinear drift in the software streams (Figure 1C). This procedure ensured the temporal alignment of data streams from eye tracking (including the world camera) and EEG. We temporarily aligned both data streams for further analysis by implementing software and hardware synchronization protocols.

### Procedure and Design

The experiment was advertised through the official platforms of Osnabrück University, and participation was scheduled once eligibility was confirmed in accordance with predefined inclusion and exclusion criteria. They were instructed to arrive at a designated location and time in the city center of Osnabrück (Figure 1D) without wearing hair products or (eye) makeup. Most of the sessions were conducted by two experimenters. The day of the experiment, each participant provided written informed consent and completed an anamnesis questionnaire. We then measured the vertex (from the nasion to the inion) and the preauricular areas of the participant’s head to place and adjust the EEG headset to ensure a proper fit. Next, eye-tracking glasses were positioned, and two electrodes were placed on the participant’s right cheek, spaced by more than 1 cm (last panel, Figure 1B). Once the EEG headset was in place, we checked the Cz electrode position and adjusted it if necessary. For all recordings, we attempted to keep electrode impedances below 10 kΩ. This was possible for 67.85% of the included sessions. We tracked electrodes with higher impedances for further analyses. Once the setup was complete and the software-level synchronization protocol had been started, we initiated the session by triggering 10 hardware-level synchronization protocol artifacts, which involved turning on the LED lights in the world camera and generating an electrical impulse to the EEG system (Figure 1B; see section *Apparatus*). Participants explored the predefined area freely for 10 minutes while an experimenter discreetly followed them to ensure safety. Afterward, we performed another set of synchronization triggers. This procedure was repeated twice, resulting in 30 minutes of exploration and three sets of triggers. A final round of hardware synchronization was then conducted at the end of the session. Participants took a self-paced break of up to 10 minutes, during which the equipment was removed and preparations were made for a second identical session. In total, we collected 60 minutes of valid data and completed eight rounds of hardware synchronization triggers (Figure 1C). Subjects were allowed to complete both sessions on the same day or different days; however, most sessions were collected on the same day (N = 22). Detailed information regarding the timing of the recording can be found in Supplementary Material 1 (Procedure and Design).

### Preprocessing

#### Data Alignment

Accurate temporal precision is critical when analyzing event-locked neural activity, as even small timing inaccuracies can substantially distort observed brain responses (Luck, 2014). To accurately align the EEG and eye-tracking data, we employed both software- and hardware-based synchronization protocols (see Figure 2 for a schematic overview). For software synchronization, we used the EEG and *RESTAPI* streams from the XDF file, along with the event file exported from Pupil Cloud. The data alignment procedure for software synchronization was performed in MATLAB (The MathWorks, Natick, MA). We began by removing eye-tracking events related to hardware synchronization triggers. Next, we trimmed the eye-tracking data to match the EEG recording window, discarding any samples recorded before the EEG start or after its end. This allowed us to establish a common reference point (i.e., the beginning of the EEG recording) for both data streams. Then, we normalized all timestamps by subtracting the time of the first sample.

**Figure 2.**
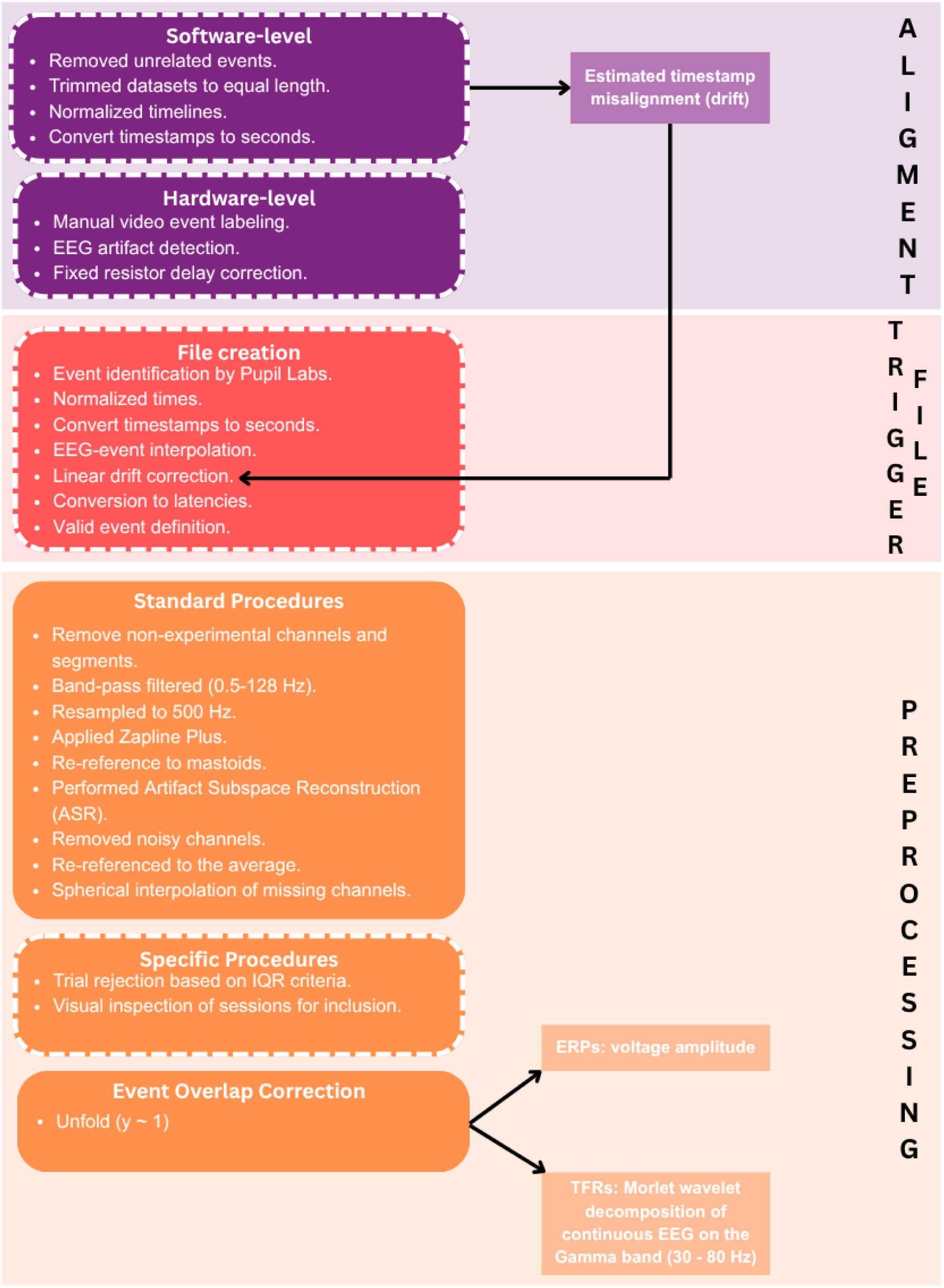
Schematic representation of the preprocessing procedure. Representation of the preprocessing pipeline implemented for the project. The schema describes the three main processes (i.e., data alignment, trigger file creation, and data preprocessing) with their respective steps in chronological order. Boxes with dotted lines indicate aspects that were specifically developed for our project, while boxes without lines represent common or standard procedures.

To quantify temporal misalignment, we computed residuals between the EEG and eye-tracking event timestamps. A linear regression model was fitted to these residuals, and the intercept was extracted. Using the fitted values, we estimated linear drift by subtracting the initial from the final predicted value. This procedure provided us with a drift estimate for each recording session using software synchronization. Across all sessions, the mean drift was −0.565 ms (SD = 3.291, min = −8.68, max = 4.858) over the complete recording period. The session-specific drift and intercept values were subsequently used to generate trigger files for each session (see *Trigger File* subsection in *Preprocessing*).

For hardware synchronization, we first manually labeled the initial video frame in the Pupil Cloud world video, where the LED lights (i.e., artifacts in the eye-tracking data) were visible. Then we exported the corresponding event file and loaded it into MATLAB along with the EEG data.

To identify the EEG artifacts generated by the impulse of the external electrodes (positioned on the participant’s right cheek), we applied a simple preprocessing pipeline on a copy of the original dataset. First, all non-EEG channels were removed, followed by bandpass filtering with a high-pass filter of 10 Hz and a low-pass filter of 128 Hz. Since the artifact was more prominent in the right-side electrodes (R1, R2, R3, and R4) and central channel (Cz, C3, and C4), we summed the signals across these channels to create a composite artifact signal.

Next, we trimmed the eye-tracking event data to align with the EEG recording window, excluding samples outside the EEG session. The event timestamps were normalized by subtracting the time of the first sample and converted to seconds. We identified the events marking the onset of the LED lights (i.e., labeled on the world video) and converted them into EEG latencies to use as temporal reference points. For each reference point, we defined a search window of ±300 ms. We applied baseline correction to the EEG signal during that window by subtracting the mean activity within it. To locate the artifact’s peak, we used the *findpeaks* MATLAB function to detect the point of maximal positive activity within the window. Each peak was marked as a hardware synchronization event in the EEG data, distinguished by blocks (A, B, C, and D). We then visually inspected the data to check for the positioning of each trigger and correct any false identifications. This procedure enabled a precise alignment between the EEG and eye-tracking timelines.

In a separate script, we imported the hardware-triggered events identified in the EEG data and converted them to seconds. To correct for the hardware-induced drift (i.e., caused by the resistor in the voltage regulator), we subtracted a fixed delay (mean = 1.122 ms). We computed the residuals between the hardware-triggered timestamps in the EEG data and the corresponding events in the eye-tracking data across all participant sessions. This analysis showed a mean hardware drift of 22 ms (SD = 23 ms, min = −186 ms, max = 81.3 ms) per session, averaged across all sessions and subjects. The small hardware drift value confirmed the accuracy of the software synchronization protocol. Overall, the small hardware and negligible software drift values indicate an accurate alignment between EEG and eye-tracking data streams, supporting further data analysis.

#### EEG Trigger File

For each EEG dataset, we generated trigger files using identified event types (saccade onset and offset, and blink onset and offset) as references. For simplicity, we refer to fixation-onset triggers as saccade-offset triggers.

Specifically, we relied on Pupil Lab’s output to identify key eye movements (i.e., blinks, fixations, and saccades). To align the eye-tracking data with the EEG, we trimmed any samples outside the established EEG recording window, as outlined in the data alignment step (see *Data Alignment* subsection in *Preprocessing*). All blink, fixation, and saccade timestamps were normalized by subtracting the first event and converted to seconds, then rounded to three decimal places to match the precision of the EEG timestamps. Then, we created an interpolated timeline spanning the EEG recording’s start and end times and mapped the fixations, saccades, and blink events by index.

To account for software-based misalignment, we linearly added the drift correction (the intercept from the previously described linear regression model) to the interpolated timeline. We extracted time points corresponding to the target event type and converted them to EEG latencies. All fixations and saccades identified as occurring during a blink, or within 10 ms before or after a blink (Dar et al., 2021), were marked as invalid. Additionally, all fixations preceded by another fixation and all saccades followed immediately by another saccade were marked as invalid. Finally, we matched all saccade events to their corresponding fixation; isolated saccades and fixations were set as invalid events.

In the trigger file, we included information about the event type (i.e., saccade onset and offset, and blink onset and offset), their latencies (in EEG samples), start times, and durations. We also included information about the validity of the saccades and fixations.

#### EEG Data

EEG data were preprocessed using a customized script, adapted from Schmidt and Nolte (2024) for the current project. Preprocessing was conducted in MATLAB, utilizing plugins from the EEGLAB toolbox (Brunner et al., 2013; Delorme & Makeig, 2004).

We converted the datasets to EEGLAB format, with double precision maintained throughout all preprocessing steps (Bigdely-Shamlo et al., 2015). Then, we removed channels that did not correspond to EEG data and non-experimental portions of data (e.g., hardware synchronization triggers). We applied a high-pass filter (0.5 Hz) and a low-pass filter (128 Hz). Following the recommendations of Klug and Kloosterman (2022), we resampled the dataset to 500 Hz and applied Zapline Plus (de Cheveigné, 2020; Klug & Kloosterman, 2022) to attenuate line noise (spectral peaks around 50 Hz). We resampled the dataset to the mastoid electrodes (L4 and R4) to establish a consistent baseline.

We implemented the *clean_rawdata* function to automatically clean noisy channels (mean = 3.4, SD = 1.5, min = 2; max = 6) and segments (mean = 96.9, SD = 91.9; min = 2, max = 330) (Kothe, 2014). Given the high noise from walking and head movements, we applied a conservative burst criterion of 20, corresponding to the standard-deviation threshold for burst removal via Artifact Subspace Reconstruction (ASR). Following noisy channel removal, the data was again re-referenced to the average. Missing channels were interpolated using spherical interpolation.

The visual inspection of the datasets still revealed significant high-frequency noise, most likely caused by walking and neck movements. Therefore, we applied a trial-rejection procedure, inspired by Ladouce and collaborators (2024) (Supplementary Material 1, *EEG Data*). Using a 1 s sliding window with 500 ms overlap, we computed window-wise IQRs and defined outlier thresholds as the median IQR ± 3.5 × the IQR of the window-wise values. Windows exceeding these thresholds in any channel were removed across all channels. After preprocessing, two experimenters (AGC and DN) independently evaluated all sessions based on inter-trial coherence and ERP morphology using conservative criteria, reaching 92.8% agreement (23 included, 7 excluded, 9 uncertain). Disagreements (N = 3) or uncertain cases were resolved by a third experimenter (PK). Subjects were generally either consistently accepted or rejected across sessions, supporting the reliability of the rejection procedure.

### Analyses

We examined single-subject ERP waveforms from −500 to 700 ms to assess differences between saccade onset and offset, and between blink onset and offset. For this, we followed a similar approach to that described by Amme and colleagues (2024): we sorted saccade-offset trials for each subject by the duration of the preceding saccade and similarly sorted blink-offset trials by the corresponding blink duration. We explored the results for every session included in the study.

Next, to correct for the effect of overlapping events, we used the Unfold toolbox (Ehinger & Dimigen, 2018). We modeled all events of interest (i.e., saccade onset and offset, and blink onset and offset) using a simple intercept model γ ∼ 1. For the time-series domain, we used the amplitude (voltage) values, while for time-frequency analyses, we computed time-frequency representations using complex Morlet wavelet convolution on continuous EEG data (Cohen, 2014). Power was estimated in the gamma range (30–80 Hz) using 20 logarithmically spaced frequencies. Wavelets were defined with 7 cycles to balance temporal and frequency resolution. The convolution was performed using the Fast Fourier Transform (FFT), and edge artifacts were removed after inverse transformation. Power was calculated as the squared magnitude of the complex signal and normalized to decibels (dB) relative to the mean power across the time series (Cohen, 2014). Gamma-band activity was obtained by averaging normalized power across 30–80 Hz, yielding a time-resolved gamma power estimate for each channel. Then, an overlap adjustment was applied to the temporal window between 500 ms before and 700 ms after event onset for ERPs and between 1 s before and 1 s after event onset for Time-Frequency representations (TFRs). For both, ERP and TFR analysis, we performed model selection by comparing the blink- and saccade-onset/offset event model with three different models with a single event for blinks and saccades, respectively. The timing of the single event was selected as the onset for one model, the offset for the second and the average of the onset and offset of the event of interest for the third.

Finally, we investigated differences between blink-related activity and saccade-related responses in the temporal and time-frequency domains. We computed cross-correlations between all possible pairs of ERPs and TFRs, using cosine similarity to assess waveform similarity across different event-locked activities for each session and subject. Following Chou & Hsu (2018) and Liu et al. (2020), we calculated cosine similarity by representing each time series as a vector and computing the cosine of the angle between them. Given the coefficients’ range (−1 to 1), we applied the Fisher z-transformation to each session’s coefficients. Due to our design, some subjects contributed with more than one session of data; hence, we averaged sessions for participants with multiple recordings (N = 16) and then computed the grand average across subjects. Then, we back-transformed the coefficients. Next, we designed a theoretical correlation matrix with ideal similarity results, under the assumption of our hypothesis of correspondence between onset and offset of blinks and saccades being true. We performed a Person correlation to test the cosine similarity coefficients of every subject and electrode with our template. Finally, to investigate significance, for each electrode separately, the subject-level Fisher z values were randomly sign-flipped across 10,000 permutations to generate a null distribution under the hypothesis that the mean representational similarity was zero. The observed mean Fisher z value was then compared against this null distribution to obtain a p-value.

All the code described in this article for preprocessing and data analysis is available online at https://github.com/AitanaGrasso-Cladera/OsnabrueckPlazaProject.

## Results

### Saccade and Blink Dynamics During Free Exploration

Given the characteristics of the experimental tasks, the subjects were able to freely explore the city center, allowing blink and saccade durations to vary naturally. This generated a high intersubject variability regarding the number of saccade and blink events. On average, we recorded a total of 5276.6 events (SD = 822.8) across sessions. As expected for a free viewing task, the number of saccadic events is larger (mean = 4430.1, SD = 649.7) than blinks (mean = 846.4, SD = 580.2). Regarding the events’ duration, blink durations showed a broad distribution across participants, with most events lasting between 200 and 300 ms (median = 245 ms, IQR = [242 ms, 300 ms]), followed by a gradual decrease in frequency for longer durations (Figure 3, green line). In contrast, saccade durations were substantially shorter and more narrowly distributed, with the highest frequencies of occurrence concentrated around 30–50 ms (median = 45 ms, IQR = [35 ms, 105 ms]) and progressively fewer long-duration events, showing a long tail (Figure 3, blue line). The distributions were largely consistent across participants, with the group-level averages highlighting the typical temporal dynamics of each event type. Importantly, the variability of durations is in the same order of magnitude as the duration of the P100 ERP component (Luck, 2014), which, being an early component, was a component of interest for the current study. This underscores that the variation in saccade and blink durations is large enough to investigate the relevance of event onset and offset to physiological signatures.

**Figure 3.**
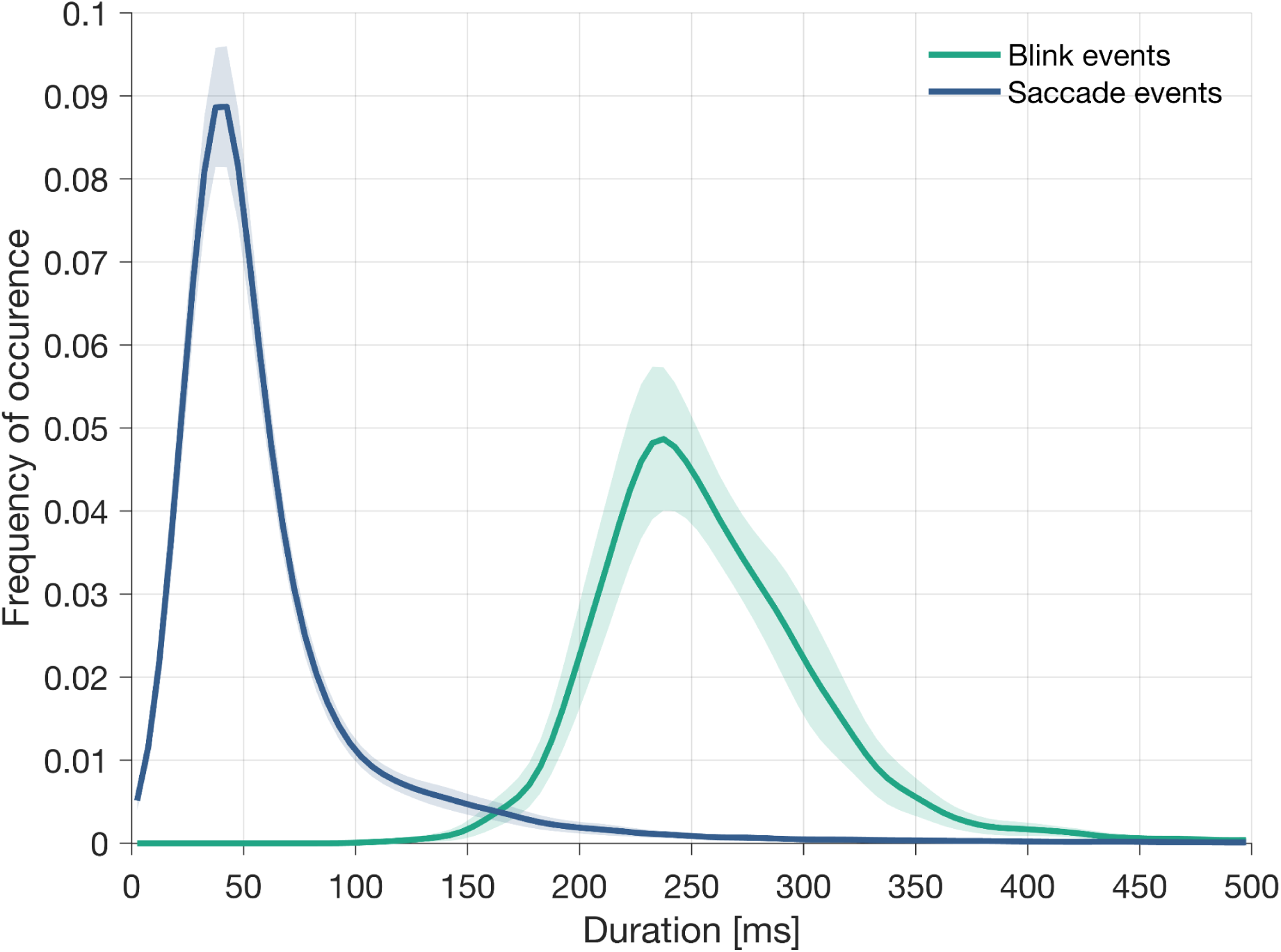
Averaged distribution of blink and saccade durations. The figure displays the averaged distribution of blink (green line) and saccade (dark blue line) durations across all sessions. Shaded areas represent 95% confidence intervals.

### Contrasting Saccade- and Blink-Offset ERPs

A subsequent step in systematically studying differences between events onset and offset, is to replicate the procedures used by Amme et al. (2024) and Nolte et al. (2025) and extend this investigation to the interruption of the visual stream caused by blinks. To address this objective, we examined single-session data and directly compared saccade-onset and saccade-offset ERPs, as well as blink-onset and blink-offset ERPs. For illustration, we selected four representative sessions from different participants (Figure 4). The data shown correspond to electrode Cz. Following the aforementioned studies, we sorted trials aligned on saccade-offset (black line; Figure 4A) for each subject by the duration of the preceding saccade (distance from white dotted line to black line; Figure 4A) to visually compare ERPs locked to saccade onset and saccade offset. We found that some structures appear to follow saccade onsets, while other visual structures seem to be better described as being locked to saccade offset. Then, we repeated the same procedure for blinks. We sorted trials aligned to blink-offset (black line; Figure 4B) by their corresponding duration (distance between white dotted line to black line; Figure 4B). Our results show that the early activity aligns partly with blink onset and partly with blink offset in varying proportions. Overall, replicating and extending previous findings, early saccadic- and blink-related responses varied across subjects, aligning with onset or offset events, or with both, with early saccadic visual responses being better aligned with saccade onset.

**Figure 4.**
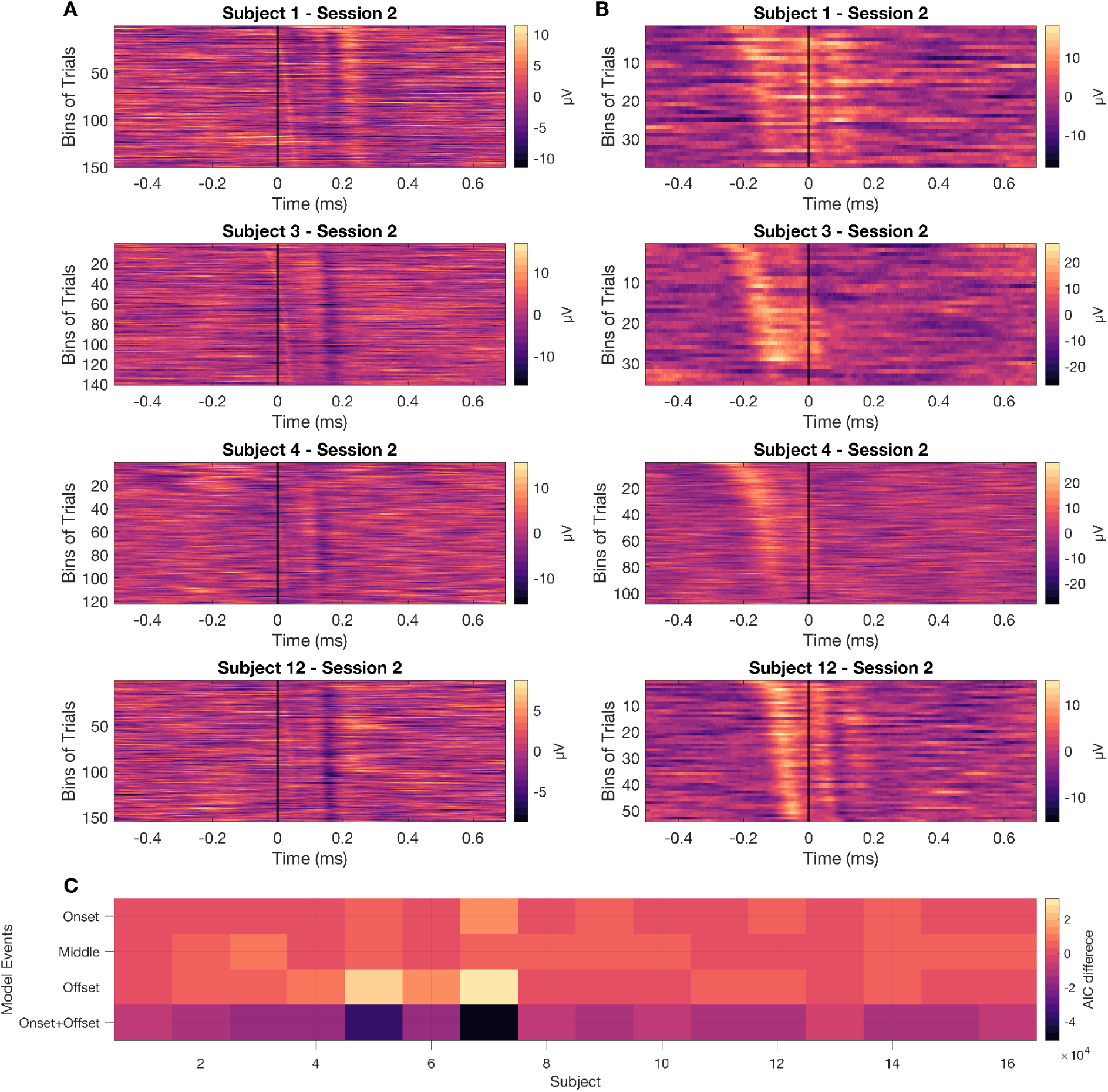
Saccade- and blink-offset ERPs at Cz, sorted by saccade and blink duration, respectively, for individual participants. (A) The four panels present all saccade offset-locked (fixation-onset) trials at electrode Cz, from single sessions of different subjects, sorted by the duration of the preceding saccade. For visualization purposes, we have binned the data (bin size = 30 trials). The top section of the *y*-axis presents the largest saccade durations, while the bottom presents the smallest durations. The vertical black line denotes the time for saccade offset (fixation onset). The white dotted line represents the saccade onset time for each trial. (B) The four panels show all blink-offset locked trials at electrode Cz. The figure shows examples from single sessions of different subjects. The trials are sorted according to the duration of the corresponding blink. For visualization purposes, we have binned the data (bin size = 15 trials). The top section of the *y*-axis presents the largest blink durations, while the bottom presents the smallest durations. The vertical black line denotes the time for the blink offset. The white dotted line represents the blink-onset time for each trial. In these figures, lighter colors represent positive amplitudes, while darker colors represent negative activations. (C) Model comparison results based on Akaike Information Criterion (AIC) values. We compared models of onset, middle latency, offset, and onset+offset. For subjects with more than one session, we averaged across sessions. In the figure, we averaged the AIC values across all models for each subject, and then subtracted it from the AIC values of each model. Lower values (darker colors) represent a better fit to the data. All subjects had negative differences for the model with onset and offset.

### Comparing Saccade- and Blink-Locked ERPs

We complement the relevance of investigating both onsets and offsets of saccadic and blink events by disentangling their contributions based on deconvolution techniques. For that purpose, we used the Unfold toolbox (Ehinger & Dimigen, 2018), which models continuous EEG data as the linear combination of overlapping event-related kernels. This deconvolution approach allowed us to correct for the temporal overlap between saccade-and blink-related activity, given their close temporal proximity, and to obtain less-biased estimates of onset- and offset-related neural responses. All kernels resulting from the deconvolution explain substantial variance of the recorded signals.

We assessed whether modeling both the onset and offset of events explains more variance in the data than modeling the onset, middle, and offset of saccades and blinks, respectively. To do this, we computed the Akaike Information Criterion (AIC) for all models. Then, we computed 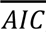, which reflects the average AIC value across all models. Across subjects, the AIC difference between the model incorporating onsets and offsets and the average (AIC_onset+offset_ - 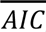) was always negative (mean = −14498, SD = 12244 [−50964, −2624]), indicating that modeling the beginning and end of the events generated lower AIC values than other models (Figure 4C). The average AIC difference for the second-best model (AIC_onset_ - 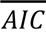) was 3545.8 (SD = 20261). In contrast, the model with only offset events presented the poorest performance, with an average AIC difference (AIC_offset_ - 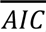) of 7625.2 (SD = 89807). The results of the model comparison favored including onset and offset events over all other models.

The deconvolution results show similarities among the different event-locked ERPs across channels (Figure 5A; Cz as an example). We observe a correspondence in the patterns of activity for saccade-onset and blink-onset ERPs, which vary in amplitude but follow a similar waveform morphology. Using root-mean-square (RMS) as a metric over the coefficients of a kernel, the blink-onset kernel was 2.49 times larger than the saccade-onset kernel. Moreover, we observe corresponding patterns in the saccade-offset and blink-offset evoked potentials. The RMS ratio is 2.3, with the blink offset kernel larger than the saccade offset kernel. Interestingly, saccade and blink onset ERPs show an early negative peak around 50 ms after event onset, followed by an increase in amplitude with positive peaks at around 100 and 200 ms. Similarly, saccade offset and blink offset ERPs show a similar pattern with an increase in amplitude around ∼150 ms before event onset, followed by a decrease after event onset. Furthermore, blink-onset and blink-offset waves exhibit opposite polarities in amplitude. Taken together, our results show that both event onset and offset contribute to ERPs for saccades and blinks, with the blink kernel, on average, more than twice as large as the saccade kernels, yet with comparable morphologies.

**Figure 5.**
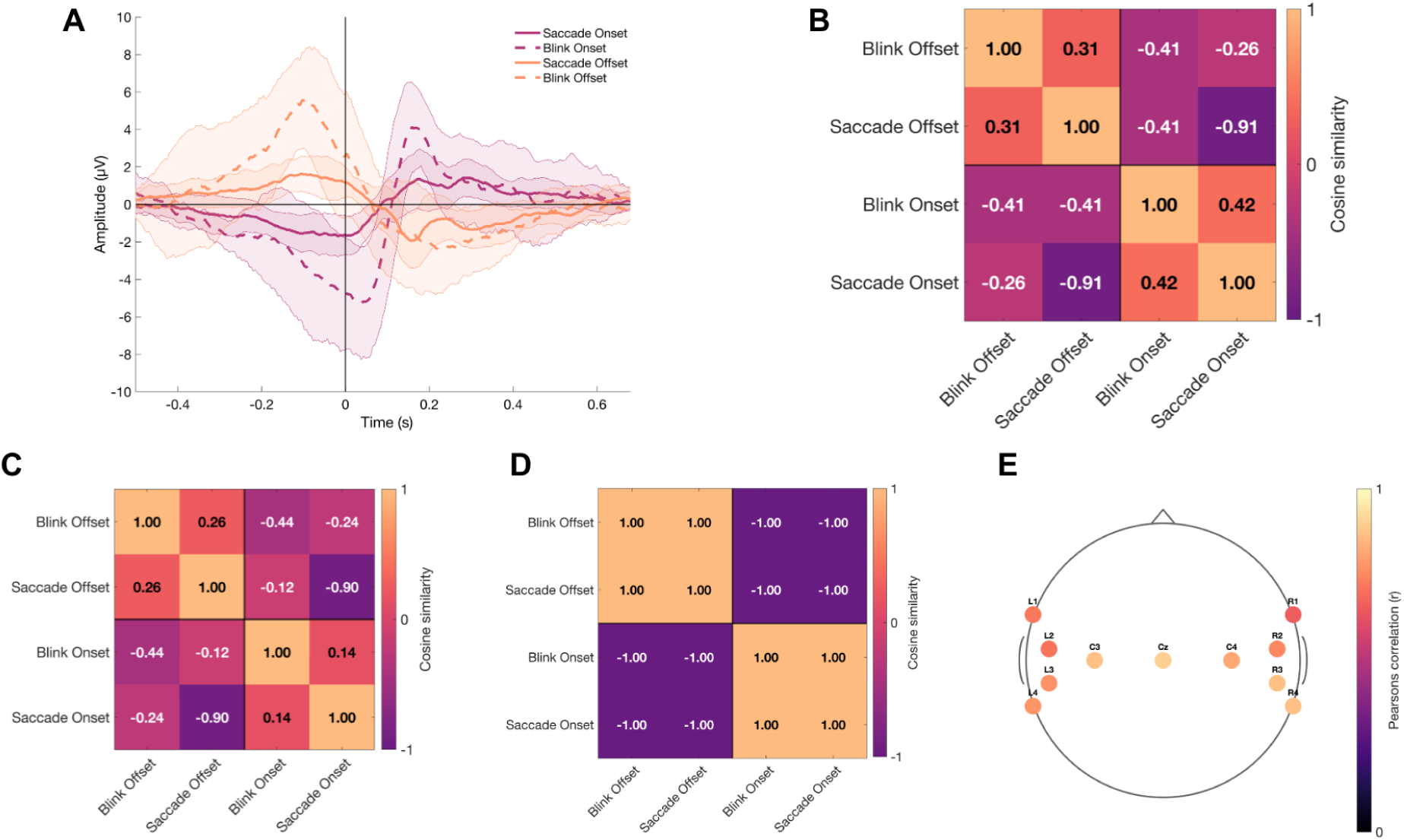
Deconvoluted potentials for all events of interest and cross-correlation results. (A) presents an example of the deconvolution results (i.e., ERPs) for the target events (i.e., blink onset and offset, and saccade onset and saccade offset), at Cz. For visualization purposes, we averaged the deconvoluted betas from subjects with multiple sessions and then computed the grand average across subjects. The black vertical line indicates the time of event onset. The continuous magenta line represents saccade onset, while the dotted magenta line represents blink onset. Similarly, the continuous orange line indicates activity locked to saccade offset, and the dotted orange line represents the blink offset. Each ERP is presented with the respective confidence interval (95%). (B) shows the results of the cross-correlation analysis, which quantifies the similarities between cortical activity locked to the different eye events of interest. Bright colors in the orange range represent positive coefficients, indicating that the compared ERPs are alike. Dark colors in the purple scale portray negative coefficients, suggesting larger differences between the compared ERPs. (C) shows the results of the cross-correlation analysis for an additional electrode, L3. (D) displays a template of a theoretical cross-correlation where blink onset and saccade onset, and blink offset and saccade offset, are identical. (E) shows Pearson’s correlations for each electrode’s cross-correlation analysis, with the template cross-correlation analysis shown in (D). The results are organized according to the EEG headset’s layout.

As a next step, we quantified the previously described similarities and differences in morphologies of the kernels. A cross-correlation analysis was performed using cosine similarity (Chou & Hsu, 2018; Liu et al., 2020) for all ERP types at time zero, without temporal shift of the kernels. The resulting cross-correlation matrix shows a block structure (Figure 5B) that supports the observed differences. Saccade onset and blink onset are positively correlated, with a cosine similarity coefficient of 0.4. Similarly, saccade offset and blink offset are positively correlated, with a coefficient of 0.3. However, saccade onset vs blink offset, saccade offset vs blink onset, blink onset vs blink offset, and saccade onset vs saccade offset showed negative similarity coefficients, with the latter combination being the most dissimilar (similarity coefficient −0.9). Overall, the similarity coefficients reflect the pattern of similarities and differences observed across the ERP kernel morphologies.

Finally, we quantify the consistency of the similarity block structure by performing a response similarity analysis. Specifically, we computed the Pearson correlation between each electrode’s correlation matrix (as an example see electrode L3 in Figure 5C) and the theoretical matrix (Figure 5D). The resulting correlations were consistently high across electrodes, though slightly smaller effects due to the increased noise levels (see Figure 5E). Correlation coefficients ranged from 0.7 (electrode R1) to 0.9 (Cz) (Figure 5E). Statistical significance of this correlation was tested by randomly sign-flipping the values across 10000 permutations using participant-level Fisher z-transformed correlation values. The results showed a significant response similarity analysis for all electrodes (p < 0.001). Taken together, our results in the time-series domain indicate a close correspondence between blink-onset and saccade-onset, as well as blink-offset and saccade-offset events.

### Comparing Saccade- and Blink-Locked Events in the Time-Frequency Domain

We replicated the same analysis for the time-frequency domain to identify differences or similarities in the Gamma band between blink- and saccade-related activity. For this, we computed Time-Frequency representations (TFRs) using complex Morlet wavelets (30–80 Hz, 20 logarithmically spaced frequencies, 7 cycles) with FFT-based convolution. The power values were normalized to decibels (dB) relative to the mean power and averaged across frequencies to obtain time-resolved gamma activity per channel. As with the time-domain analysis, this deconvolution technique allowed us to assess onset- and offset-related neural activity. All kernels resulting from the deconvolution explain substantial variance of the recorded signals.

To assess whether onset and offset events should be modeled separately, we investigated the variance explained by our model with four events and compared it to models with only two events, representing (the beginning, middle, or end of) saccades and blinks. To do this, we computed the Akaike Information Criterion (AIC) for all models and then calculated 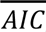 which reflects the average AIC value across all models. Across subjects, the AIC difference between models was indicative that modeling the onset and offset of events generated lower AIC values than modeling the time of any of the other events (mean = −88818; SD = 128240; [-512024, 2904]). This result was consistent for 14 of 16 subjects, for the two remaining subjects the model with the middle event outperformed the model with beginning and end. The average AIC difference with the second-best model (AIC_middle_ - 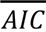) was 8776, with a SD of 44738. In contrast, the model with only offset events presented the poorest performance, with an average AIC difference (AIC_offset_ - 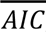) of 59417 (SD = 53558). The results of the model comparison for the time-frequency analysis favored the modeling of onset and offset events.

The deconvolution results show similarities among the different event-locked TFRs across channels (Figure 6A; Cz as an example). Comparing the kernels for blink onset and saccade onset revealed an RMS over time of 3.3, indicating that the blink-onset kernel is larger than the saccade-onset one. This demonstrates that saccade and blink onsets have similar amplitudes and morphologies, though a slight temporal shift can be observed. Furthermore, we see a similar pattern for saccade-offset vs. blink-offset: the TFRs have similar amplitudes and morphologies over time with a RMS ratio over time of 0.9, highlighting a larger amplitude for the blink offset kernel in comparison to the saccade offset kernel. In particular, it is possible to see a clear difference between the pairs of events mentioned earlier, characterized by positive (saccade offset and blink offset) and negative (saccade onset and blink onset) deflections before event onset. This shows a similar canceling effect around 0 dB as described for the time-series domain. Shortly after event onset (∼0 to ∼150 ms), saccade onset, and saccade offset TFRs present a clear and sharp peak, which is less defined for blink onset and offset. Finally, we see a convergence of the TFRs towards the end of the trial (from ∼450 ms to ∼850 ms). In line with the time-series domain, the time-frequency results support the observation of event onset and offset contributions to the TRFs for both saccades and blinks.

**Figure 6.**
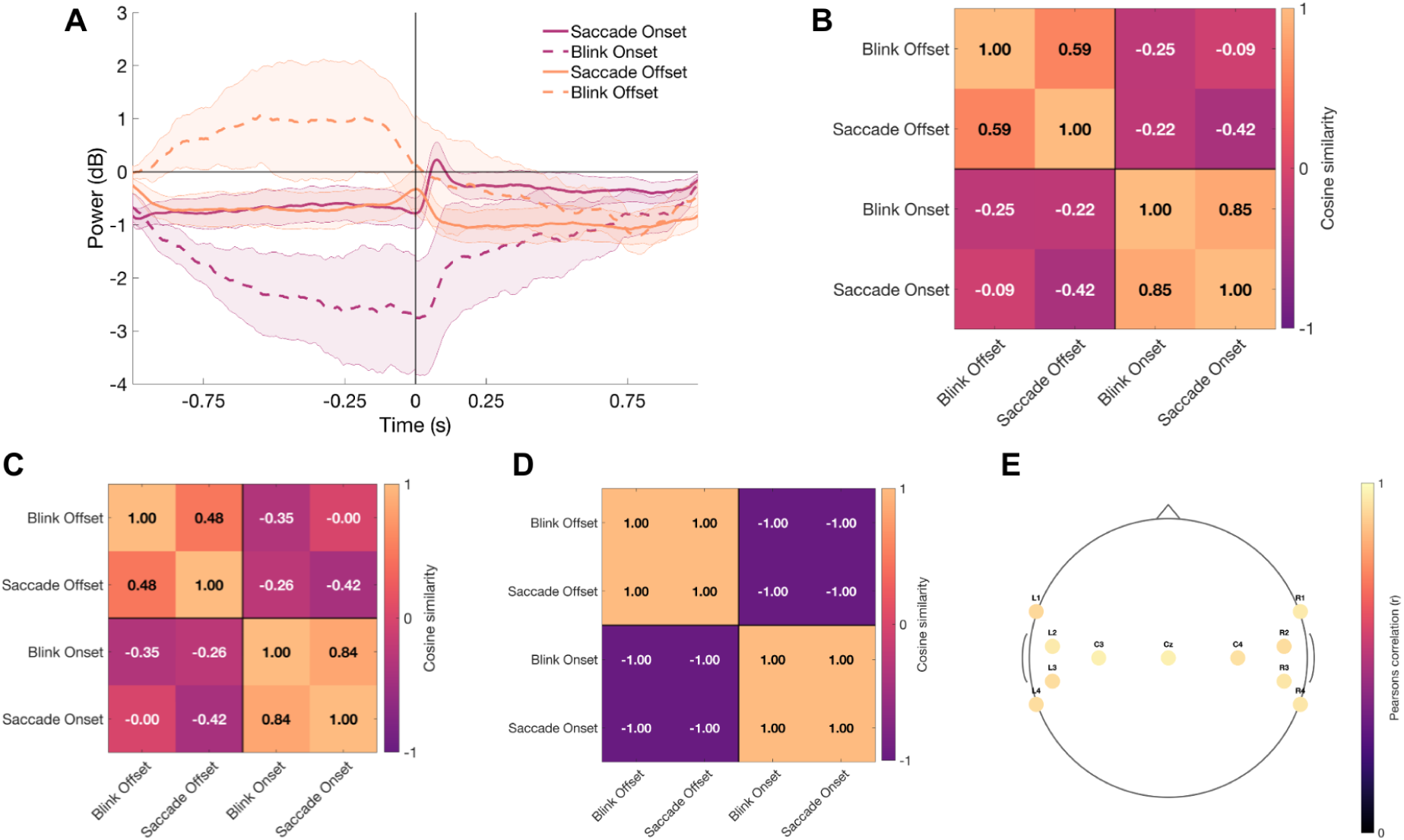
Deconvoluted Time-Frequency representations (TFRs) for all events of interest and cross-correlation results. (A) presents an example of the deconvolution results (i.e., TF) for the target events (i.e., blink onset and offset, and saccade and saccade offset), at Cz. For visualization purposes, we averaged the deconvoluted betas from subjects with multiple sessions and then computed the grand average across subjects. The black vertical line indicates the time of event onset. The continuous magenta line represents saccade onset, while the dotted magenta line represents blink onset. Similarly, the continuous orange line indicates activity locked to saccade offset, and the dotted orange line represents the blink offset. Each TFR is presented with the respective confidence interval (95%). (B) shows the results of the cross-correlation analysis, which quantifies the similarities between cortical activity locked to the different eye events of interest. Bright colors in the orange range represent positive coefficients, indicating that the compared TFRs are alike. Dark colors in the purple scale portray negative coefficients, suggesting larger differences between the compared TFRs. (C) shows the results of the cross-correlation analysis for an additional electrode, L3. (D) displays a template of a theoretical cross-correlation where blink onset and saccade onset, and blink offset and saccade offset, are identical. (E) shows Pearson’s correlations for each electrode’s cross-correlation analysis, with the template cross-correlation analysis shown in (D). The results are organized according to the EEG headset’s layout.

Following the same procedure as for ERPs, we quantified similarities using a cross-correlation analysis with cosine similarity (Chou & Hsu, 2018; Liu et al., 2020). The correlation was computed at time zero, without any temporal shift of the kernels. For TFRs, saccade vs blink onset presented a cosine similarity coefficient of 0.9, while saccade offset vs blink offset presented a coefficient of 0.6 (Figure 6B). Conversely, saccade onset vs blink offset, saccade offset vs blink onset, blink onset vs blink offset, and saccade onset vs saccade offset showed lower similarity coefficients. TRF comparisons underscore similarities for onset events, as well as offset events, for blinks and saccades.

As with the time domain analysis, we systematically investigated the consistency of the cosine-correlation block structure across all electrodes by performing a response similarity analysis. For this, we constructed a matrix representing the theoretical results for saccades and blinks, assuming an identical pattern (Figure 6D), and then computed Pearson’s correlation between each electrode’s matrix (as an example see electrode L3 in Figure 6C) and this theoretical matrix. These results were corroborated by the remaining electrodes, which showed the same pattern as Cz (Figure 6E). Notably, correlating all electrodes with an ideal matrix revealed correlation coefficients above 0.9, indicating a systematic effect across electrodes. Statistical significance of this correlation was tested by randomly sign-flipping the values across 10000 permutations using participant-level Fisher z-transformed correlation values. The results showed a significant response similarity to the theoretical matrix for all electrodes (p < 0.001). Taken together, our time-frequency results support the time-series findings, indicating a close correspondence between blink-onset and saccade-onset, as well as between blink-offset and saccade-offset events.

## Discussion

In this study, we investigated the real-world neural dynamics of naturally occurring self generated eye events, saccades and blinks. Our findings successfully replicate recent evidence that saccade-related activity is robustly driven by event onset rather than offset alone (Amme et al., 2024; Nolte et al., 2025), while critically extending this principle to blinks by demonstrating a comparable onset and offset division. Quantifying this observation using deconvolution techniques, we demonstrated that the underlying neural kernels are highly correlated when comparing saccade onset to blink onset, as well as saccade offset to blink offset. Additionally, the results in the time-frequency domain aligned with these observations, providing further evidence of the correspondence between neural signatures at blink onset and saccade onset, and at blink offset and saccade offset. Together, these findings highlight the relevance of saccade and blink onset and offset in structuring ongoing visual input during natural free viewing and emphasize the correspondence between the neural dynamics of these events.

The observed similarities between saccade- and blink-onset events and between saccade- and blink-offset events may reflect overlapping mechanisms involved in active visual sampling and oculomotor control. The shared activity could, for example, correspond to activity in a common central oculomotor network, including regions such as the frontal eye fields and the inferior precentral sulcus, two areas implicated in the generation of saccades and blinks (Kato & Miyauchi, 2003). Further, both events have been associated with the reallocation of attention (Deubel & Schneider, 1996; Nakano et al., 2013). The similarities between blink- and saccade-onset activity may therefore reflect shared mechanisms for preparing for self-generated changes in the visual input, potentially related to predictive processing of active free viewing (Crapse & Sommer, 2008; Friston, 2010). Relatedly, blinks and saccades are accompanied by a suppression of visual processing, reducing perceptual input during self-generated motion or visual occlusion (Burr, 2005; Johns et al., 2009; Ridder & Tomlinson, 1997; Willett et al., 2023). Under such an account, our results could be interpreted as shared processes of visual suppression, potentially due to internal predictions of sensory consequences which may contribute to perceptual stability during active free vision (Crapse & Sommer, 2008; Friston, 2010). Complementing this suppression, saccade and blink offset events may correspond to the processing of novel visual input, potentially resembling stimulus onset in controlled laboratory paradigms, although previous work has also noted differences between these processes (Amme et al., 2024; Ladouce et al., 2022). Overall, these similar neural signatures suggest that blinks and saccades share underlying mechanisms for managing sensory input and maintaining visual continuity during free vision.

Our experiment employed a methodological framework that leverages naturally occurring eye events, identified using eye-tracking technologies, as temporal anchors for studying neural dynamics during unconstrained behavior. Specifically, we examined EEG activity associated with self-generated saccades and blinks, extending previous MoBI and free-viewing approaches that have used ocular events to investigate event-related neural responses in naturalistic settings (Alyan et al., 2023; Ladouce et al., 2022; Wascher et al., 2022; Wunderlich & Gramann, 2021). Importantly, in these free-viewing approaches, the ocular events are subject driven, and therefore are not independent variables, requiring the use of deconvolution techniques (Ehinger & Dimigen, 2018). A particular strength of our approach is that these events were identified directly from eye-tracking recordings rather than inferred from EEG signals. Although the combination of EEG and eye tracking is well established, eye-tracking information is most commonly used to improve EEG preprocessing by facilitating the detection and correction of ocular artifacts (Klug & Kloosterman, 2022; Mannan et al., 2016; Plöchl et al., 2012). In contrast, studies investigating blink- and saccade-related potentials often identify ocular events from EEG-derived signals, such as independent component activations or vertical electrooculogram (vEOG) channels (Alyan et al., 2023; Wascher et al., 2022; Wunderlich & Gramann, 2021). However, previous work has demonstrated that integrating eye tracking with EEG improves the detection and removal of ocular artifacts and preserves a greater proportion of neural activity than EEG-only approaches (Mannan et al., 2016; Plöchl et al., 2012), including under free-viewing conditions (Dimigen, 2020). As a result, the use of eye-tracking–based event identification in our free-viewing study provides a principled framework for investigating the neural correlates of ocular behavior in naturalistic environments.

There are some challenges and limitations that have to be mentioned. One of the main limitations concerned the EEG setup, which consisted of an 11-channel system only, primarily covering the midline and temporal regions. Together with the high level of noise inherent to mobile EEG recordings, this led us to focus our analyses predominantly on the Cz electrode. The limited spatial resolution of this setup prevented reliable topographic analyses and source localization. Nevertheless, although the electrode montage was not optimal for investigating activity in early visual areas, the consistency of the results across electrodes suggests that the limited occipital coverage may not have had a major influence on the main findings of the present study. Another challenge was the increased susceptibility of mobile recordings to noise and recording instabilities, which often result in elevated participant exclusion or dropout rates (Debener et al., 2015; Plöchl et al., 2012). This was also the case in our study, reducing the number of usable datasets and potentially affecting statistical power and reproducibility. Finally, the number of recorded events differed considerably across event types, with substantially fewer blinks than saccades and fixations. Such imbalances are common in MoBI paradigms, where behavioral events cannot be experimentally controlled to occur at equivalent rates, thereby challenging comparisons across conditions (Nolte et al., 2025). Despite these limitations, the present study demonstrates that meaningful and reliable estimates of fixation-, saccade-, and blink-related neural activity can be obtained from mobile EEG recordings in naturalistic settings, supporting the feasibility of investigating event-related brain dynamics during real-world behavior.

This project systematically characterized neural signatures of saccade and blink-onset and -offset events in a naturalistic, free-viewing setting. The naturalistic setting allowed us to investigate the dynamics of these events as they naturally occur without task-related influences that could alter them (Bachurina & Arsalidou, 2022; Bentivoglio et al., 1997; Gandhi, 2012; McMonnies, 2020). However, future studies may need to design more controlled experiments to further explore their relevance and relationship. If the neuronal activity related to blink- and saccade-onset has similar patterns, the observed activity should not relate to the specific mechanisms triggering saccades or blinks individually, but to a more general consequence of transiently interrupting visual input. Paradigms explicitly investigating these eye events and their related visual input, such as event-contingent change-blindness paradigms (Henderson & Hollingworth, 2003), may therefore help disentangle the underlying mechanisms. To support such investigations, neuroimaging techniques with higher spatial resolution, such as MEG or fMRI, could provide a detailed characterization of the cortical networks involved. Future studies may also benefit from further refining the timing of event onsets and offsets. Given that eye-movement classification algorithms differ in their accuracy in identifying different eye events, improving the precision and consistency of identification may be useful (Nir & Deouell, 2026). Overall, our findings on the shared neural dynamics of saccades and blinks warrant experimental follow-up approaches that isolate the underlying mechanisms and uncover their contributions to natural perception.

## Acknowledgements

We would like to thank Isabelle Sander for her contribution to developing the initial code for exploring the hardware-level synchronization protocol. We would also like to thank Aiko-Theres Dubrall for testing the hardware setup for the experiment, conducting the first pilot sessions, and contributing to data collection. Furthermore, we would like to thank Fabia Segatz for her support during data collection. Finally, we would like to extend our gratitude to Moritz Querdel for developing and testing the hardware synchronization device, as well as for his extensive participation as a pilot subject of the complete experimental hardware setup. We would like to thank the German Research Foundation DFG which supported this project in the context of funding the Research Training Group “Situated Cognition” (GRK 274877981).

## Funding

This project is supported by the Federal Ministry for Economic Affairs and Energy (BMWE) on the basis of a decision by the German Bundestag (grant reference number KK5743901RL4). This project is also funded by the Deutsche Forschungsgemeinschaft (DFG, German Research Foundation – Projektnummer GRK 274877981). Furthermore, this project is supported by the ERC Starting Grant “TIME” (grant number 101039524). Aziz Muhammed Akkaya was funded by ERASMUS+ and Ibn Haldun University during his stay for this project.

## Notes

https://github.com/AitanaGrasso-Cladera/OsnabrueckPlazaProject

https://osf.io/dvgue/overview

### Competing Interest Statement

Aitana Grasso-Cladera reports equipment, drugs, or supplies was provided by mBrainTrain.

