## Supplementary Material 1 for "Exploring Neural Signatures of Self-Initiated Saccadic and Blink Events in the Real World"

### Procedure and Design

The experiment was advertised through the official platforms of the University. Interested individuals were contacted and scheduled for participation after their eligibility was confirmed in accordance with predefined inclusion and exclusion criteria. Before their session, participants received an online overview of the experimental procedure. They were instructed to arrive at a designated location and time in the city center (Figure 1D), without wearing hair products or makeup. Two experimenters conducted most of the sessions.

The day of the experiment, each participant provided written informed consent and completed an anamnesis questionnaire. We then measured the vertex (from the nasion to the inion) and the preauricular areas of the participant’s head to place and adjust the EEG headset to ensure a proper fit. Next, eye-tracking glasses were positioned, and two electrodes were placed on the participant’s right cheek, spaced approximately by more than 1cm (last panel, Figure 1B). Once the EEG headset was in place, we checked the Cz electrode position and adjusted it if necessary. For all recordings, we attempted to keep electrode impedances below 10 kΩ. This was possible for 67.85% of the included sessions. We kept track of electrodes with higher impedances for further analyses.

Once the setup was complete and the software-level synchronization protocol had been started, we initiated the session by triggering 10 hardware-level synchronization protocol artifacts, which involved turning on the LED lights in the world camera and generating an electrical impulse to the EEG system (Figure 1B). Then, participants were instructed to walk at a normal pace within a designated area of the city, simulating the experience of waiting for a friend, while avoiding interactions with objects and pedestrians and entering stores. These instructions mirrored those used by Nolte et al. (2025) in a virtual environment. Participants explored the area freely for 10 minutes while being discreetly followed by an experimenter for safety reasons. Afterward, we performed another set of synchronization triggers. This procedure was repeated twice, resulting in 30 minutes of exploration and three sets of triggers. A final round of hardware synchronization was then conducted at the end of the session. Participants took a self-paced break of up to 10 minutes, during which the equipment was removed, and preparations were made for a second identical session. In total, we collected 60 minutes of valid data and completed eight rounds of hardware synchronization triggers (Figure 1C). Subjects were allowed to complete both sessions on the same day or different days; however, most sessions were collected on the same day (N = 22).

To maintain consistency between participants, we restricted our data collection to Monday through Saturday during typical working hours (9:30 AM to 7:30 PM), when pedestrian traffic was relatively stable. Furthermore, to avoid different sunlight conditions, we standardized session timing to one or two time windows per day, depending on the sun's position. We did not record data during rainy days.

### Validation of the Eye-Tracking Data

We computed the effective sampling rate by computing the residual between timestamps for the video (mean = 29.98 Hz, SD = 0.00, min = 29.97, max = 29.99) and the eye data (mean = 198.18 Hz, SD = 0.833, min = 195.06, max = 199.68) across all recordings. These results demonstrate that the effective sampling rate aligned with expectations, even when synchronizing between multiple devices. We then conducted a visual inspection to evaluate the accuracy of ocular event detection (e.g., fixations and saccades) provided by the Pupil Labs built-in algorithm. This involved examining eye-position data (X and Y coordinates) over time, the detected onset and offset times of fixations and saccades, and the durations of saccadic events. Our visual inspection supported the algorithm's accuracy. Consequently, we could rely on the ocular event markers generated by Pupil Cloud without requiring a secondary detection algorithm.

### EEG Data

The visual inspection of the datasets still revealed significant high-frequency noise, most likely caused by walking and neck movements. Consequently, we applied a procedure to remove trials with aberrant activity inspired by the work of Ladouce and collaborators (2024), who used the same EEG system. For this, we first implemented a 1s sliding window with a 500 ms overlap and computed the IQR for each window. We calculated the median of the window-wise IQRs and the IQR of these values (from now on, the *general IQR*). We defined upper and lower thresholds as the median IQR ± 3.5 × *general IQR*. Then, we computed the IQR for each channel in each window and compared it against the thresholds. If a window exceeded either threshold at any channel, it was marked as an outlier and removed from all channels. After preprocessing was completed, two experimenters (AGC and DN) independently rated all sessions from all subjects for inclusion, using a conservative approach. We implemented a visual inspection of inter-trial coherence and assessed ERP shapes. Overall, we had 92.8% agreement on the selection (23 included, 7 excluded, 9 uncertain). A third experimenter (PK) acted as a mediator in discussions about the final decision when the two experimenters were uncertain or disagreed (N = 3). Coincidentally, the subjects kept were those with already one good session, while the subjects rejected were generally rejected in both sessions. This resulted in most subjects having both sessions excluded or included, and gave us confidence that the rejection protocol worked well.
